# Phylogenetic tree-aware positive-unlabeled deep metric learning for phage–host interaction identification

**DOI:** 10.64898/2025.12.31.696981

**Authors:** Yao-zhong Zhang, Boschung Tobias, Seiya Imoto

## Abstract

Phages are viruses that infect bacteria and play essential roles in shaping microbial communities. Identifying phage–host interactions (PHIs) is crucial for understanding infection dynamics and developing phage-based therapeutic strategies. Recent deep learning approaches have shown great promise for PHI prediction; however, their performance remains constrained by the limited number of experimentally validated positive pairs and the overwhelming abundance of unlabeled or non-validated samples. Moreover, most existing models overlook higher-level phylogenetic relationships among hosts, which could provide valuable structural priors for guiding representation learning.

To address these challenges, we propose a phylogenetic tree–aware positive–unlabeled deep metric learning framework for phage–host interaction (PHI) identification. Unlike traditional approaches that train classification models to strictly separate positive and negative phage–host pairs, the proposed method learns representations under supervision from both confirmed positive PHIs and host phylogenetic tree constraints on non-positive samples. The proposed method can seamlessly formalize contrastive learning and deep metric learning within the same framework that explicitly optimizes PHI encoders with biological constraints in the learning functions. We show that this metric learning formulation outperforms conventional contrastive learning approaches that enforce separation between positive and negative samples without consistently aligning the learned representations with evolutionary distances. Experiments on the Cherry benchmark dataset and metagenome Hi-C multi-host dataset demonstrate that our approach enhances species-level prediction accuracy, improves cross-host generalization, and yields more interpretable representations of phage–host relationships.

## 1 Introduction

Bacteriophages (phages) are viruses that infect bacteria and archaea, playing central roles in microbial ecosystems by modulating bacterial populations, driving horizontal gene transfer, and shaping community structure and function. Accurate identification of phage–host interactions (PHIs) is therefore critical for understanding infection dynamics and for applications such as phage therapy, microbiome engineering, and antimicrobial design. Yet experimental PHI mapping is laborious and biased toward cultivable strains, leaving most phage–host relationships uncharacterized.

Computational approaches offer a promising complement to experimental phage–host interaction (PHI) discovery. Classical methods primarily rely on explicit biological signals, including sequence alignment and compositional similarity (e.g., k-mer profiles), CRISPR spacer matches, and ecological co-occurrence patterns. To improve robustness and coverage, recent ensemble frameworks integrate multiple heterogeneous signals within unified pipelines, exemplified by iPHoP [1]. Deep learning–based approaches further advance PHI prediction by learning non-linear mappings between viral and host features. Depending on the modeling formulation, representative methods can be broadly categorized into link-prediction-based and classification-based frameworks. For link-prediction–based frameworks, phage–phage and phage–host relationships are jointly represented as a heterogeneous graph, on which graph-based encoders such as graph convolutional networks (GCNs) or graph attention networks (GATs) are applied to learn node representations for interaction prediction [2, 3, 4]. For classification-based frameworks, features derived from representations at the nucleotide and protein levels, either separately or jointly, as well as receptor-binding protein (RBP) representations, are used as inputs to deep learning classifiers to predict target hosts or binary infection labels [5, 6, 7]. Besides the above formalization approaches, either as classification or link prediction, representation learning through contrastive learning has been applied to learn phage-host interaction encoders that make phages and infected hosts close in the new embedding space [8]. More recently, advances in genomic and protein language models have been leveraged for PHI prediction by extracting informative sequence representations from pretrained models [7, 9].

Despite these advances, existing deep models are often limited by the scarcity of experimentally validated phage–host interactions. On one hand, most positive pairs correspond to single-host associations, whereas the number of candidate negative hosts is substantially larger, leading to severe label imbalance in classification-based training. On the other hand, non-positive pairs may include both truly non-infecting and unverified interactions, making the naive treatment of all non-positive hosts as negatives potentially incorrect. Recent large-scale ecological analyses reveal that phages with broad host ranges are common across ecosystems [10], highlighting the incompleteness and ambiguity of current phage–host interaction labels. Based on these observations, we propose a novel PHI learning framework that integrates phylogenetic guidance for unlabeled samples from the perspective of positive–unlabeled learning. Specifically, based on deep metric learning, we formulate a unified cross-entropy framework that includes InfoNCE objectives for positive phage–host pairs from supervised labels and cross-entropy objectives for negative pairs with phylogenetic distance guidance. The underlying objective is to make the learned phage and host representations align with phylogenetic distances—that is, hosts that are phylogenetically closer (e.g., within the same genus) are expected to exhibit more similar or related infection (or distance) patterns, which are used as additional general constraints for negative samples.

We evaluated the proposed method on the Cherry and Hi-C metagenome multi-host datasets. The proposed method learns phylogeny-aware host representations by incorporating host phylogenetic relationships during training. Compared with the margin-based contrastive learning method, the proposed method not only significantly accelerates the overall training speed but also improves species-level prediction accuracy. Compared with other commonly used PHI methods, the proposed method also demonstrates better prediction accuracy for multi-host predictions.

## 2 Method

### 2.1 Overall framework

In this work, we formulate phage–host interaction (PHI) identification within the framework of *positive–unlabeled* (PU) deep metric learning. Different from conventional PU learning approaches, which typically operate only on phage representations while treating hosts as categorical labels, our method embeds both phages and hosts into a shared representation space. Infection relationships are then inferred based on distances between their embeddings in this space.

Given a set of phages 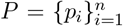 and candidate hosts 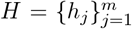, we learn phage and host encoders Enc_*p*_ and Enc_*h*_ that map genomes into a common embedding space. Here, we use a shared encoder such that Enc_*p*_ = Enc_*h*_. Let **Z**_*p*_ = Enc_*p*_(*P*) ∈ ℝ^*n*×^ and **Z**_*h*_ = Enc_*h*_(*H*) ∈ ℝ^*m*×*D*^. Pairwise similarity score (logit) between a phage *p*_*i*_ and a host *h*_*j*_ is defined as

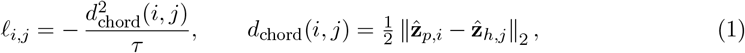

where 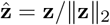 denotes *ℓ*_2_-normalized embeddings and *τ* > 0 is a temperature parameter.

#### PHI identification with infoNCE

PHI identification is inherently a positive–unlabeled (PU) problem, where experimentally validated phage–host pairs serve as positives and all other pairs remain unlabeled. InfoNCE is often applied in this setting by implicitly assuming unlabeled pairs to be negatives. With similarity scores defined in Eq. (1), the standard multi-positive InfoNCE loss is given by

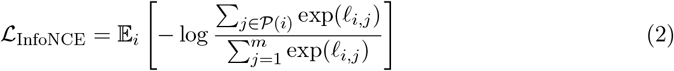

where 𝒫 (*i*) denotes the set of known infecting hosts for phage *p*_*i*_. This contrastive objective encourages each phage to align closely with its infecting hosts while uniformly repelling all unlabeled hosts. However, such uniform treatment of negatives ignores biological structure among candidate hosts and prevents the incorporation of phylogenetic information.

Recent works have explored principled extensions of standard InfoNCE that relax the implicit assumption that all unlabeled samples are true negatives. Acharya *et al*. [11] formulate contrastive learning explicitly under a positive–unlabeled (PU) setting and correct the bias of InfoNCE by accounting for hidden positives within the unlabeled pool, thereby avoiding uniform repulsion of all non-positive samples. From a complementary perspective, Robinson *et al*. [12] demonstrate that the effectiveness of contrastive learning critically depends on how negatives are weighted, and propose hardness-aware negative sampling schemes that emphasize informative (hard) negatives instead of treating all negatives equally. Together, these works suggest that the uniform treatment of unlabeled samples in Eq. (2) is suboptimal: unlabeled instances differ in both semantic relevance and reliability, and should therefore contribute unequally to the contrastive objective.

Motivated by these insights, one possible refinement is to incorporate phylogenetic structure into the treatment of negative samples by defining a Laplacian tree kernel based on phylogenetic distances, which can be used to modulate the hardness of unlabeled hosts. In conventional contrastive learning formulations, similarity is defined purely by the distance between embedded phage and host representations (e.g., derived from FCGR features), and optimization proceeds by simultaneously pulling positive pairs together while pushing all non-positive pairs apart. However, this objective can be difficult to optimize in practice. On the one hand, the model is required to enforce a clear separation between positive and negative samples; on the other hand, for hosts with highly similar FCGR representations—such as those belonging to the same genus—it is challenging to further separate them more strongly than other less similar hosts without destabilizing training.

To address this issue, we hypothesize that learning could be more stable and easier when both positive and negative samples are optimized along a consistent direction, rather than through the opposing attractive–repulsive forces inherent to standard contrastive objectives. Based on this perspective, we propose a unified cross-entropy (CE) formulation that reinterprets InfoNCE as a special case of cross-entropy with hard (one-hot) labels, and extends it by assigning structured, soft target distributions to unlabeled hosts. This formulation naturally enables the integration of phylogenetic information into the learning objective, allowing biologically related hosts to be repelled in a graded manner rather than uniformly. It preserves the basic framework of contrastive learning, while providing a principled mechanism for incorporating biological constraints into phage–host representation learning.

#### From infoNCE to cross-entropy formulation with soft labels

We develop CE4PHI, which formulates a positive–unlabeled (PU) deep metric learning objective to learn phage–host relationships using a tree-guided cross-entropy loss. For a given phage *p*_*i*_, the loss is defined as

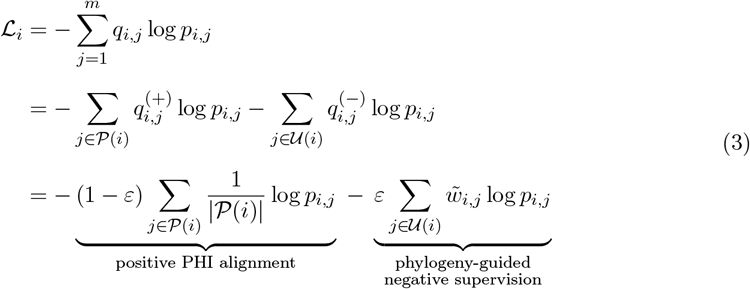

Here, the model-induced interaction probability is defined via a softmax over distance-based logits,

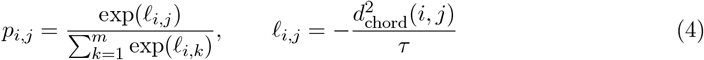

Positive probability mass is assigned uniformly across known infecting hosts, while the remaining mass *ε* is distributed among unlabeled hosts according to their phylogenetic proximity,

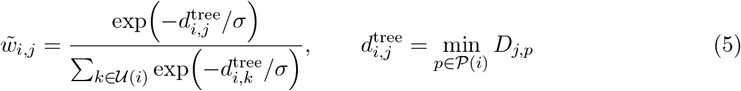

where *D* is the normalized host phylogenetic distance matrix and *σ* > 0 controls the decay rate.

This formulation yields a unified deep metric learning objective in which positives are assigned a higher probability mass while unlabeled hosts receive smoothly decaying mass according to their evolutionary distance. By reshaping the embedding geometry through probability reallocation rather than explicit contrastive repulsion, the proposed loss enables biologically informed, same-direction optimization for multi-host PHI identification.

### 2.2 Building phylogenetic tree for bacterial host

The phylogenetic distances among hosts were derived from the Genome Taxonomy Database (GTDB) [13]. For the provided host assemblies or metagenome-assembled genomes (MAGs), we used GTDB-Tk (v2.5.2) to reconstruct the phylogenetic relationships among bacterial host genomes. GTDB-Tk implements a standardized approach based on the Genome Taxonomy Database (GTDB), in which a set of universally conserved, single-copy marker genes (120 bacterial or 122 archaeal markers) are identified using Prodigal for gene prediction and HMMER searches against TIGRFAM/Pfam models. The identified marker proteins are aligned individually, filtered, and concatenated into a genome-wide multiple sequence alignment. A phylogenetic tree is then inferred from the concatenated alignment using FastTree under appropriate protein substitution models. An archaeal phylum (p__Thermoproteota) was designated as the out-group in order to root the bacterial tree and a bacterial phylum ((p__Bacillota)) was used as the outgroup for the root archaea tree. For the cherry dataset, among the 223 bacterial hosts, there are 221 bacteria and 12 archaea. After building tree distance matrices for bacteria and archaea separately, we combined them by filling the cross-group entries with an additional 20% of the largest distance in the combined matrix. The generated tree distances are used for calculating 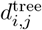

## 3 Experiments

### 3.1 Evaluation datasets

We evaluated the proposed method on CHERRY benchmark dataset and HiC metagenome data. CHERRY is a commonly used benchmark dataset [14] that contains 1940 phages and 223 bacterial hosts. Like most other existing datasets, it is a single-host dataset where each phage is assigned one infected host in the dataset. For a concise description, we use *dataset*_*CHERRY*_ as the abbreviation in the following sections. We followed their methods of splitting the datasets. Similarly, we use *dataset*_*HiC*_ as an abbreviation for the Hi-C–derived metagenomic dataset. Phage genomes were obtained from the proximity-ligation-based metagenomic Hi-C (metaHiC) dataset reconstructed by Bignaud et al. [10], together with the associated metadata tables. The data set contains 249 hosts and 2883 phages, including 487 phages with multiple hosts. The corresponding bacterial host genomes were downloaded from NCBI GenBank [15] based on species-level taxonomy. Any host with taxonomic rank below species was eliminated, leaving 52 hosts and 406 phages, of which 46 have multiple hosts. We constructed two complementary dataset splits for *dataset*_*HiC*_ to evaluate model performance. In the first split, all single-host cases were assigned to the training and validation set, while all multi-host cases were reserved exclusively for testing. This setup enables us to assess the model’s ability to generalize from single-host patterns to previously unseen multi-host scenarios. In the second split, we employed a stratified partitioning strategy based on the number of hosts, ensuring that both single- and multi-host instances are proportionally represented in the training and test sets. This configuration provides a more balanced evaluation of the model under realistic distributional conditions. The detailed data summary is shown in the supplementary material.

### 3.2 Evaluation metrics and baseline methods

Most existing PHI prediction methods are evaluated under a single-host assumption, typically using top-K accuracy at different taxonomic levels. Beyond conventional top-1 and top-K accuracy metrics, we extend the standard top-1 evaluation by introducing a *multi-host accuracy* metric, which explicitly accounts for phages with multiple known hosts. For a phage with a gold host set 𝒢_*i*_ of size *N*_𝒢__*i*_= |𝒢_*i*_|, we consider the top-*N*_𝒢__*i*_ predicted hosts 𝒫_*i*_ and compute the number of correctly recovered hosts as |𝒢_*i*_ ∩ 𝒫_*i*_ |. Aggregating over all phages, the multi-host accuracy is defined as:

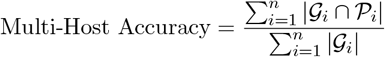

This metric generalizes the single-host top-1 accuracy to settings where each phage may naturally infect multiple hosts.

To quantify the advancements introduced in this work, we compare our method against four previously published models. First, we benchmark the proposed model against its direct predecessor, CL4PHI [8], which allows us to isolate the improvements attributable to the architectural and training modifications introduced in this work. In addition to CL4PHI, we include representative PHI prediction baselines spanning probabilistic, alignment-free, and supervised deep-learning paradigms. PHP [16] predicts hosts using a Gaussian model over k-mer frequency differences between viral and prokaryotic genomes. PHIST [17] is a fast alignment-free tool that links viruses to hosts by counting exact shared k-mers between their sequences. In addition, we report updated Cherry results obtained via the PhaBOX2 platform [2]. Rather than performing direct model-level comparisons with baseline methods, we primarily use them as software-level references to contextualize performance on the evaluated datasets.

### 3.3 CE4PHI improves species-level PHI prediction accuracy of CL4PHI

We first compare the original CL4PHI model, which employs a margin-based contrastive loss with a single positive host per phage, against CE4PHI, a tree-aware variant that replaces the margin formulation with a unified cross-entropy objective capable of incorporating all known infecting hosts simultaneously through soft-label reallocation. The comparison is conducted on the two datasets introduced in this work, *dataset*_CHERRY_ and *dataset*_HiC_.

For fairness and reproducibility, both models adopt the same parameter settings as reported in the original implementation^1^. CE4PHI follows a hyperparameter search strategy consistent with that used in CL4PHI, with detailed configurations provided in the Supplementary Material. Both models were trained under identical experimental conditions using the same random seed (e.g., 123) and hardware (Nvidia A6000 Ada GPU) for 300 epochs on *dataset*_CHERRY_ and both splits of *dataset*_HiC_, with the optimal checkpoint selected based on validation prediction accuracy. Performance is benchmarked using the previously introduced multi-host accuracy and top-*k* accuracy metrics, where top-*k* predictions are computed based on the model’s distance-based scoring function. Since *dataset*_CHERRY_ contains no multi-host cases, the multi-host accuracy metric reduces to standard top-1 accuracy for this dataset.

As shown in Table 1, CE4PHI consistently improves species-level accuracy across all evaluation cutoffs (Top-1/3/5/10: 0.6120/0.7461/0.7918/0.8849), outperforming CL4PHI(0.5946/0.7303 /0.7776/0.8580) in fine-grained host identification. At the genus level, CL4PHI achieves a slightly higher Top-1 accuracy (0.7397 vs. 0.7177), whereas CE4PHI surpasses CL4PHI at larger *K* values (Genus Top-3/5/10: 0.8517/0.8991/0.9290 vs. 0.8454/0.8659/0.9101). To further disentangle the role of negative supervision, we include an ablation variant **CE4PHI (***ε* = 0), in which the cross-entropy objective is restricted to positive samples only and no phylogenetically guided negative probability mass is assigned. While CE4PHI (*ε* = 0) yields modest improvements over CL4PHI at certain cutoffs (e.g., Species Top-5: 0.7950 vs. 0.7776), it consistently underperforms the full CE4PHI model, particularly at higher genus-level cutoffs (e.g., Genus Top-10: 0.9243 vs. 0.9290). These results confirm that explicit soft negative reallocation guided by phylogenetic distance is a key contributor to the observed performance gains.

**Table 1:** Comparison of model performance on the CHERRY dataset. CE4PHI denotes the proposed tree-aware cross-entropy formulation with soft negative supervision, while CL4PHI uses a margin-based contrastive objective.

| Model | Species Accuracy |  |  |  | Genus Accuracy |  |  |  |
| --- | --- | --- | --- | --- | --- | --- | --- | --- |
|  | Top1 | Top3 | Top5 | Top10 | Top1 | Top3 | Top5 | Top10 |
| **CL4PHI** | 0.5946 | 0.7303 | 0.7776 | 0.8580 | **0.7397** | 0.8454 | 0.8659 | 0.9101 |
| **CE4PHI** | **0.612** | **0.7461** | 0.7918 | **0.8849** | 0.7177 | **0.8517** | **0.8991** | **0.929** |
| **CE4PHI ($\varepsilon=0$)** | 0.5994 | 0.7413 | **0.795** | 0.8849 | 0.7066 | 0.847 | 0.8912 | 0.9243 |

#### 3.3.1 Margin-based and InfoNCE-style objectives induce complementary prediction behaviors

As shown in Table 1, we observe a performance trend between the two learning objectives (This is also observed on *dataset*_HiC_). Compared with CE4PHI, the original CL4PHI achieves slightly higher genus-level Top-1 accuracy, whereas CE4PHI yields superior performance at the species level as well as for larger *K* in Top-*K* evaluation. These differences are not incidental, but instead reflect fundamental distinctions in how margin-based and InfoNCE-style objectives shape the geometry of the embedding space.

CL4PHI adopts a margin-based contrastive loss that explicitly enforces a minimum distance between positive and negative pairs. Once the margin constraint is satisfied, the corresponding gradients rapidly diminish, leading to sparse updates for well-separated samples. As a result, the optimization is dominated by boundary enforcement rather than by continuous refinement of relative distances. In the PHI setting, where hosts within the same genus are inherently similar in sequence space, this objective naturally encourages phage embeddings to collapse toward a dominant genus-level cluster while pushing other genera beyond the decision boundary. Such block-wise separation favors coarse-grained discrimination and directly benefits genus-level Top-1 prediction. However, because the margin constraint does not impose further penalties once satisfied, the internal structure among species within the same genus is only weakly constrained.

In contrast, CE4PHI with *ε* = 0 corresponds to a multi-positive InfoNCE-style cross-entropy objective. Rather than relying on an explicit distance threshold, this formulation normalizes similarity scores over all candidate hosts through a softmax operation, ensuring that every negative sample contributes to the gradient throughout training. This probability-driven optimization emphasizes global ranking over local boundary conditions, continuously encouraging the model to resolve fine-grained differences among closely related hosts. Consequently, multiple species within the same genus compete for probability mass, leading to a more structured and discriminative intra-genus geometry in the embedding space. While this property improves species-level resolution and Top-*K* performance, it may also distribute probability mass across related genera, making CE4PHI less aggressive in winner-takes-all genus-level Top-1 decisions. The observed performance trade-offs highlight the distinct geometric inductive biases of the two objectives. Margin-based contrastive learning prioritizes stable, coarse decision boundaries, whereas InfoNCE-style cross-entropy favors continuous, fine-grained embedding refinement.

#### 3.3.2 CE4PHI learns phylogeny-aware host representations

Figure 2 compares the t-SNE visualizations of joint phage–host embeddings derived from the original flattened FCGR representation, the margin-based CL4PHI encoder, and the proposed CE4PHI encoder on the CHERRY dataset. In the FCGR space, hosts and phages exhibit extensive overlap with weak geometric organization, indicating that compositional similarity alone is insufficient to recover phylogenetically meaningful structure.

**Figure 1:**
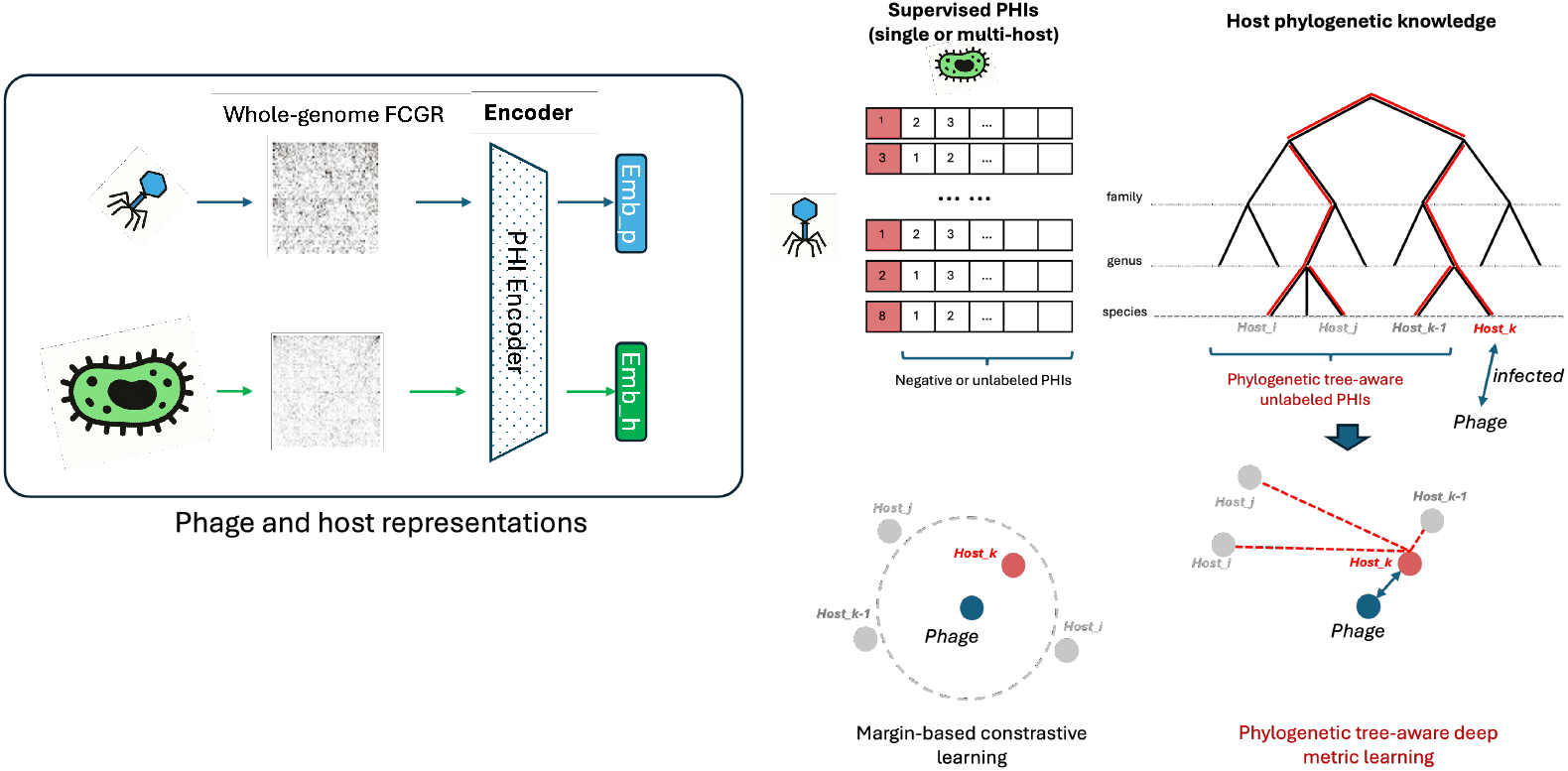
Overall pipeline of the phylogenetic tree–aware positive–unlabeled deep metric learning framework for PHI identification. Phage and host genomes are first represented using two-dimensional k-mer (frequency chaos game representations) as the raw input. An encoder is then trained to model phage–host infection relationships using experimentally validated positive PHI labels. Rather than treating all non-positive candidate hosts as equally negative, the proposed framework incorporates host phylogenetic relationships into the deep metric learning objective, allowing the encoder to account for evolutionary proximity among hosts during training.

**Figure 2:**
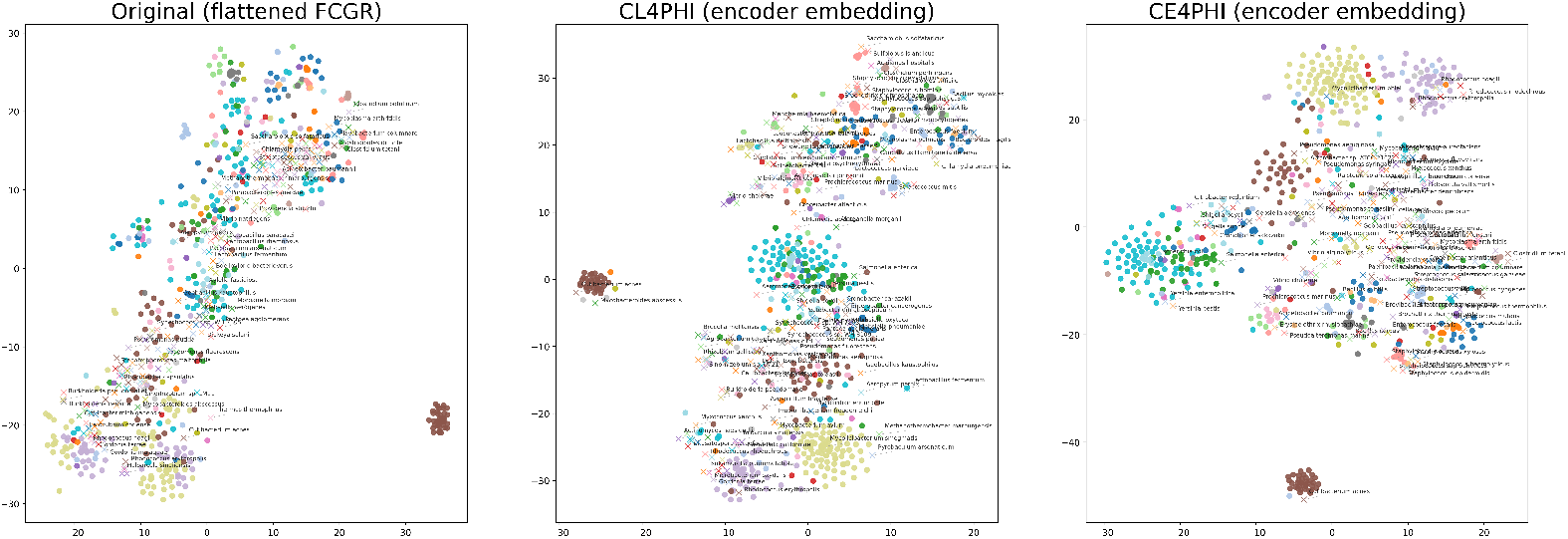
t-SNE visualization of joint phage–host embeddings on the CHERRY dataset, showing the original flattened FCGR space (left), CL4PHI encoder embeddings (middle), and CE4PHI encoder embeddings (right). Phages are represented as points (•), while host genomes are indicated by crosses (×). CE4PHI yields a more structured organization consistent with host phylogeny, preserving intragenus separability while maintaining global taxonomic coherence.

Applying the CL4PHI encoder introduces coarse clustering effects, suggesting that contrastive learning improves global separability. However, hosts belonging to the same genus frequently collapse into overly compact regions, and boundaries between closely related genera remain blurred. This behavior is particularly evident for genera such as *Streptococcus* and members of the *Enterobacteriaceae*, reflecting the limitations of margin-based objectives that enforce uniform repulsion among all non-positive hosts without accounting for phylogenetic relatedness.

In contrast, the CE4PHI embedding displays a markedly more structured geometry aligned with host phylogeny. Closely related genera within the *Enterobacteriaceae*, including *Escherichia, Shigella, Salmonella, Yersinia*, and *Cronobacter*, are embedded in adjacent yet distinguishable regions, preserving their known evolutionary relationships while avoiding representation collapse. In particular, *Escherichia* and *Shigella* appear in close proximity, whereas *Salmonella* and *Yersinia* occupy nearby but separable subregions, consistent with their phylogenetic distances. Similarly, *Streptococcus* species form a coherent genus-level cluster in the CE4PHI embedding, within which individual species remain separable, demonstrating that fine-grained intra-genus variation is preserved rather than compressed. Comparable improvements are observed for other Firmicutes such as *Clostridium*, whose species form compact yet internally structured clusters that are clearly separated from other genera. We note that a small number of hosts exhibit less well-defined separation. In particular, archaeal genera such as *Acidianus* and *Saccharolobus* show partial overlap with neighboring regions, which likely reflects data imbalance and the limited representation of archaeal hosts in the CHERRY dataset, as well as intrinsic limitations of k-mer–based FCGR features for extremophilic genomes.

Overall, these visualizations demonstrate that CE4PHI learns an embedding space that preserves fine-grained intra-genus structure while maintaining global phylogenetic consistency. The resulting geometry provides a clear visual explanation for the improved species-level resolution observed on the CHERRY dataset, as quantitatively reported in Table 1.

#### 3.3.3 Effect of the soft negative mass parameter *ε*

In CE4PHI, the parameter *ε* explicitly controls the allocation of probability mass between confirmed positive hosts and unlabeled candidate hosts in the cross-entropy objective. For each phage *p*_*i*_, a fraction 1 − *ε* of the target mass is assigned to experimentally validated host(s), while the remaining fraction *ε* is distributed across unlabeled hosts according to their phylogenetic proximity. When *ε* = 0, the objective reduces to a hard contrastive formulation equivalent to standard InfoNCE, whereas increasing *ε* progressively introduces phylogeny-guided soft supervision from unlabeled hosts.

Figure 3 shows the effect of *ε* on species- and genus-level top-*K* accuracy. We observe a clear unimodal trend across all metrics: performance improves when a small amount of soft negative mass is introduced (*ε* ≈ 0.02–0.05), remains stable within a narrow intermediate range, and gradually degrades as *ε* increases further. In the small-*ε* regime, soft negative supervision acts as a regularizer, reducing overconfident single-host assignments while preserving fine-grained discrimination among closely related hosts.

**Figure 3:**
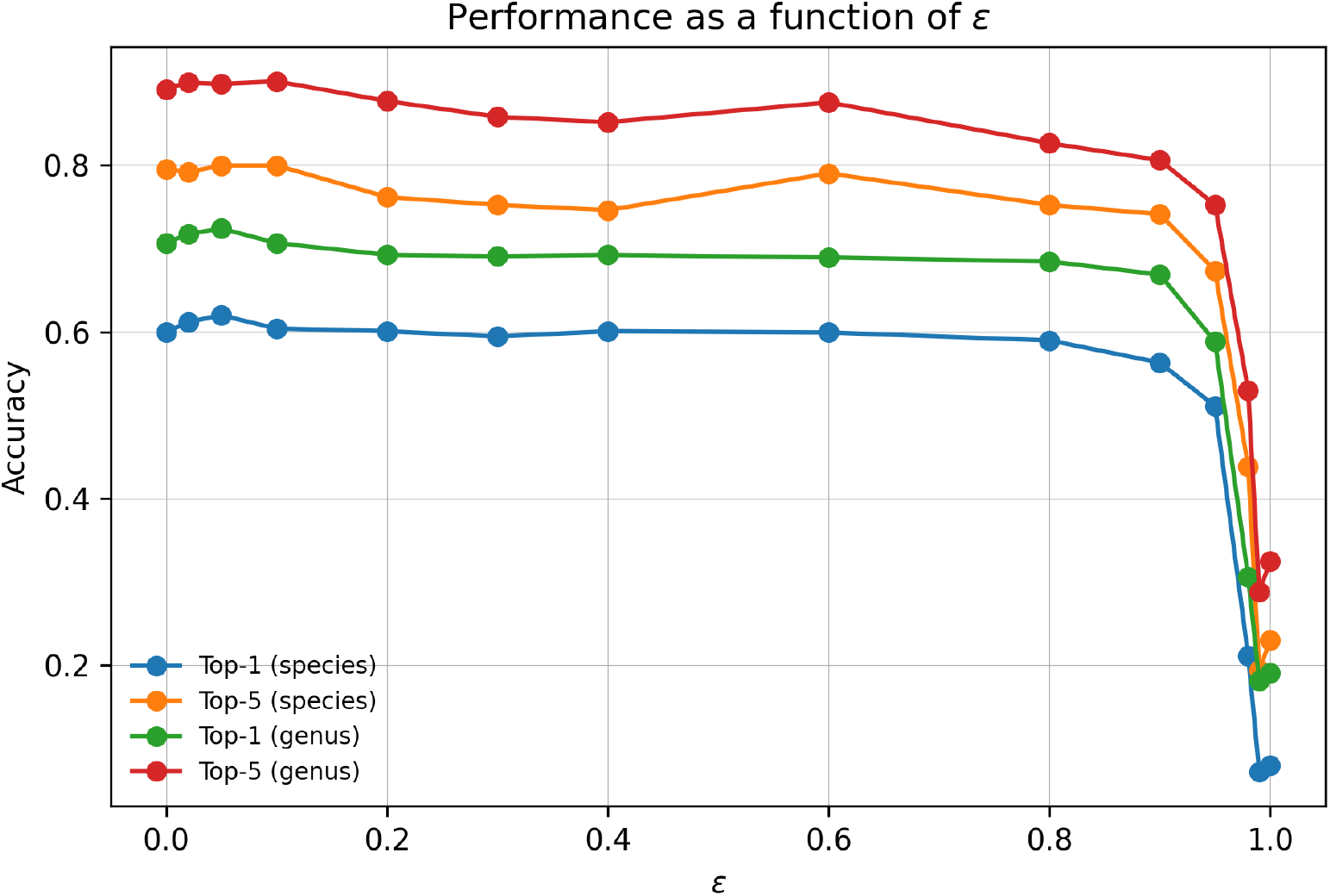
Effect of the tree-aware cross-entropy weight *ε* on prediction performance on the CHERRY dataset. The figure shows species-level and genus-level top-*K* accuracy as a function of *ε*.

Notably, performance does not collapse immediately when *ε* > 0.5, but instead remains relatively stable until *ε* becomes very large. This behavior can be attributed to the large number of candidate hosts in the CHERRY benchmark (223 hosts), over which the unlabeled mass *ε* is normalized and distributed. As a result, each individual unlabeled host receives only a small fraction of supervision, and the optimization process remains anchored by the positive term as long as 1 − *ε* is non-negligible. When *ε* approaches 1, the positive supervision term vanishes and the objective becomes dominated by tree-guided unlabeled supervision, effectively fitting a smooth phylogenetic prior rather than phage-specific host signals, leading to sharp performance degradation.

### 3.4 Metagenome HiC data Multi-host evaluation

We further evaluated the proposed method under two complementary evaluation settings on HiC multi-host dataset. (1) In-data evaluation, where CL4PHI models are trained and tested within the Hi-C host–interaction dataset, assessing their performance on the same data distribution. (2) Cross-data evaluation, where models are trained on the CHERRY dataset and evaluated on the metagenomic Hi-C dataset, providing a stringent test of generalization across datasets with distinct sequencing platforms, sample compositions, and host diversity. CL4PHI and CE4PHI are trained using the CHERRY split supplied by its original as in the CHERRY paper.

#### 3.4.1 In-data evaluation

We first evaluate CL4PHI and CE4PHI under the *in-data* setting on the Hi-C multi-host dataset, where training and testing are performed on the same data distribution but across different splits. Table 2 reports both multi-host accuracy and Top-*K* prediction accuracy at the species and genus levels. Across both splits, CE4PHI consistently outperforms CL4PHI in terms of multi-host accuracy. On Split 1 (Sp-1), CE4PHI improves species-level multi-host accuracy from 0.4000 to 0.5333 and genus-level accuracy from 0.8214 to 0.8393. Similarly, on Split 2 (Sp-2), CE4PHI achieves higher multi-host accuracy at both the species level (0.6224 vs. 0.5408) and the genus level (0.8471 vs. 0.8353). These gains indicate that the proposed tree-aware cross-entropy formulation is more effective at capturing multiple plausible host associations per phage than the margin-based contrastive objective. Consistent improvements are also observed in Top-*K* accuracy, particularly at the species level. On Sp-1, CE4PHI improves species Top-1 accuracy from 0.5000 to 0.5870 and achieves uniform gains at larger *K* values (Top-3/5). On Sp-2, CE4PHI yields a notable increase in species Top-1 accuracy (0.6585 vs. 0.5854), while maintaining comparable performance at higher *K*. These results suggest that CE4PHI enhances fine-grained species discrimination without sacrificing recall among multiple candidate hosts. At the genus level, both methods achieve near-saturated performance at larger *K* values, reflecting the relatively coarse resolution of genus-level prediction in the Hi-C dataset. Nevertheless, CE4PHI consistently improves genus-level Top-1 accuracy on Sp-2 (0.8780 vs. 0.8659) and yields higher Top-K accuracy across splits. This indicates that CE4PHI better ranks phylogenetically related hosts even when multiple genera are plausible.

**Table 2:**
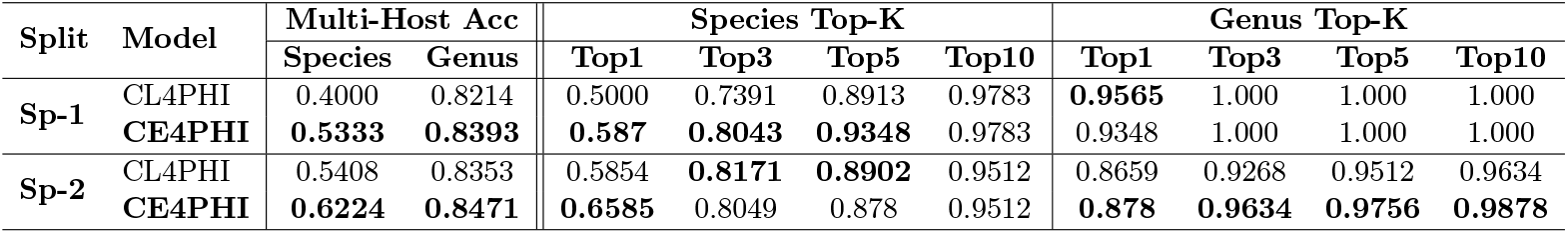
Evaluation of multi-host accuracy and top-K accuracy for models trained on *dataset*_*HiC*_. Model hyper-parameters for the dataset are determined through the 1-best accuracy of the validation set. lr=1e-5, batch size=32.

| Split | Model | Multi-Host Acc |  | Species Top-K |  |  |  | Genus Top-K |  |  |  |
| --- | --- | --- | --- | --- | --- | --- | --- | --- | --- | --- | --- |
|  |  | Species | Genus | Top1 | Top3 | Top5 | Top10 | Top1 | Top3 | Top5 | Top10 |
| Sp-1 | CL4PHI | 0.4000 | 0.8214 | 0.5000 | 0.7391 | 0.8913 | 0.9783 | **0.9565** | 1.000 | 1.000 | 1.000 |
|  | **CE4PHI** | **0.5333** | **0.8393** | **0.587** | **0.8043** | **0.9348** | 0.9783 | 0.9348 | 1.000 | 1.000 | 1.000 |
| Sp-2 | CL4PHI | 0.5408 | 0.8353 | 0.5854 | **0.8171** | **0.8902** | 0.9512 | 0.8659 | 0.9268 | 0.9512 | 0.9634 |
|  | **CE4PHI** | **0.6224** | **0.8471** | **0.6585** | 0.8049 | 0.878 | 0.9512 | **0.878** | **0.9634** | **0.9756** | **0.9878** |

#### 3.4.2 Cross-data evaluation

We then evaluate model generalization under the *cross-data* setting, where models are trained on the CHERRY dataset and tested on the metagenomic Hi-C dataset. This setting introduces substantial domain shift, including differences in sequencing protocols, sample composition, and host diversity, and therefore provides a stringent test of robustness.

As shown in Table 3, CE4PHI consistently outperforms CL4PHI across both splits in terms of multi-host accuracy. On Split 1 (Sp-1), CE4PHI improves species-level multi-host accuracy from 0.3167 to 0.4500 and genus-level accuracy from 0.5357 to 0.7857. Similar gains are observed on Split 2 (Sp-2), where CE4PHI increases species-level multi-host accuracy from 0.3061 to 0.4286 and genus-level accuracy from 0.5059 to 0.6824. These improvements indicate that the proposed tree-aware cross-entropy formulation generalizes effectively across datasets than the margin-based contrastive objective.

**Table 3:**
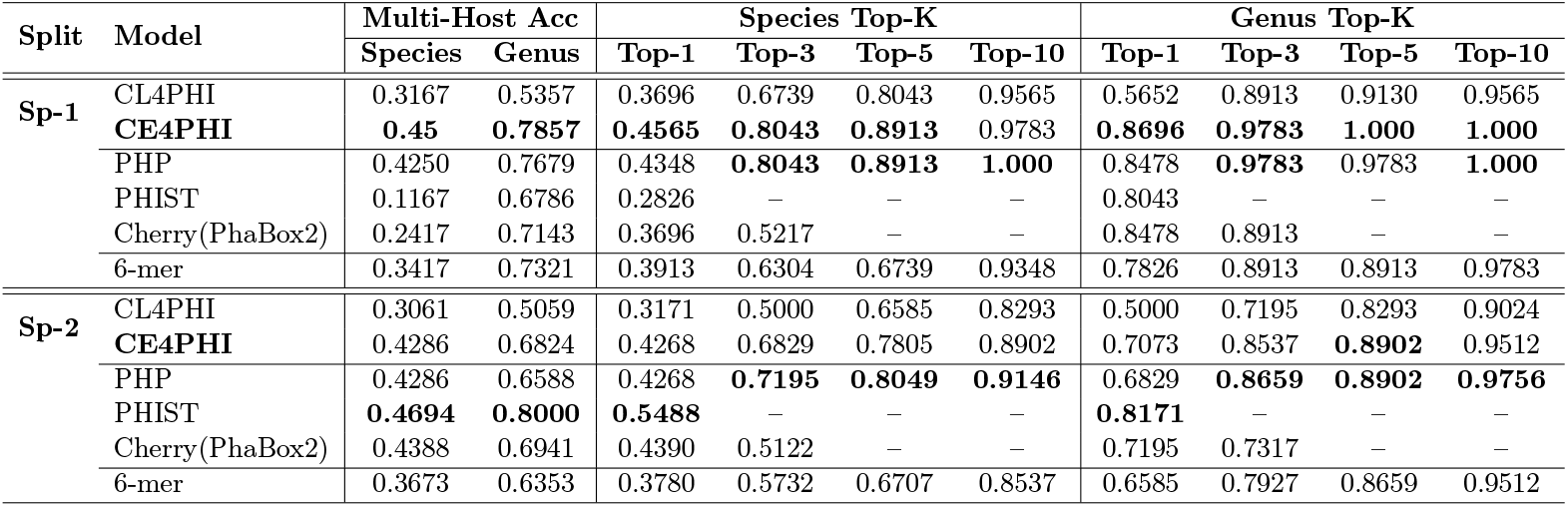
Comparison of multi-host accuracy and species/genus top-k accuracy for the cross-data evaluation on metagenome Hi-C two data splits.

| Split | Model | Multi-Host Acc |  | Species Top-K |  |  |  | Genus Top-K |  |  |  |
| --- | --- | --- | --- | --- | --- | --- | --- | --- | --- | --- | --- |
|  |  | Species | Genus | Top-1 | Top-3 | Top-5 | Top-10 | Top-1 | Top-3 | Top-5 | Top-10 |
| Sp-1 | CL4PHI | 0.3167 | 0.5357 | 0.3696 | 0.6739 | 0.8043 | 0.9565 | 0.5652 | 0.8913 | 0.9130 | 0.9565 |
|  | **CE4PHI** | **0.45** | **0.7857** | **0.4565** | **0.8043** | **0.8913** | 0.9783 | **0.8696** | **0.9783** | **1.000** | **1.000** |
|  | PHP | 0.4250 | 0.7679 | 0.4348 | **0.8043** | **0.8913** | **1.000** | 0.8478 | **0.9783** | 0.9783 | **1.000** |
|  | PHIST | 0.1167 | 0.6786 | 0.2826 | — | — | — | 0.8043 | — | — | — |
|  | Cherry(PhaBox2) | 0.2417 | 0.7143 | 0.3696 | 0.5217 | — | — | 0.8478 | 0.8913 | — | — |
|  | 6-mer | 0.3417 | 0.7321 | 0.3913 | 0.6304 | 0.6739 | 0.9348 | 0.7826 | 0.8913 | 0.8913 | 0.9783 |
| Sp-2 | CL4PHI | 0.3061 | 0.5059 | 0.3171 | 0.5000 | 0.6585 | 0.8293 | 0.5000 | 0.7195 | 0.8293 | 0.9024 |
|  | **CE4PHI** | 0.4286 | 0.6824 | 0.4268 | 0.6829 | 0.7805 | 0.8902 | 0.7073 | 0.8537 | **0.8902** | 0.9512 |
|  | PHP | 0.4286 | 0.6588 | 0.4268 | **0.7195** | **0.8049** | **0.9146** | 0.6829 | **0.8659** | **0.8902** | **0.9756** |
|  | PHIST | **0.4694** | **0.8000** | **0.5488** | — | — | — | **0.8171** | — | — | — |
|  | Cherry(PhaBox2) | 0.4388 | 0.6941 | 0.4390 | 0.5122 | — | — | 0.7195 | 0.7317 | — | — |
|  | 6-mer | 0.3673 | 0.6353 | 0.3780 | 0.5732 | 0.6707 | 0.8537 | 0.6585 | 0.7927 | 0.8659 | 0.9512 |

At the species level, CE4PHI achieves consistent improvements in Top-*K* accuracy across both splits. On Sp-1, CE4PHI improves species Top-1 accuracy from 0.3696 to 0.4565 and shows substantial gains at larger *K* values (Top-3/5/10: 0.8043/0.8913/0.9783). On Sp-2, CE4PHI similarly outperforms CL4PHI at all evaluated cutoffs (Top-1/3/5/10: 0.4268/0.6829/0.7805/0.8902 vs. 0.3171/0.5000/0.6585/0.8293), demonstrating improved ranking of correct hosts under domain shift. Genus-level performance exhibits the same trend. CE4PHI yields large improvements in genus Top-1 accuracy on both Sp-1 (0.8696 vs. 0.5652) and Sp-2 (0.7073 vs. 0.5000), while maintaining strong Top-*K* performance at larger cutoffs. These results suggest that CE4PHI more reliably preserves phylogenetic structure when transferring across datasets, particularly at coarse taxonomic resolution.

Compared with existing methods evaluated in the same cross-data setting, including PHP, PHIST, PhaBox2, and a 6-mer baseline, CE4PHI achieves competitive or superior performance across most metrics, especially in species-level Top-1 accuracy and multi-host prediction. While PHIST attains strong Top-1 accuracy on Sp-2, its evaluation is limited to Top-1 metrics and does not provide consistent performance across splits. In contrast, CE4PHI demonstrates stable improvements across all evaluated criteria, highlighting its robustness under realistic cross-dataset transfer. The cross-data evaluation confirms that incorporating phylogenetically guided soft negative supervision enhances generalization in phage–host interaction prediction. By reallocating probability mass among related hosts rather than enforcing hard inter-class margins, CE4PHI mitigates overfitting to dataset-specific host distributions and yields more reliable multi-host predictions under domain shift.

### 3.5 CE4PHI can significantly reduce training time

CE4PHI exhibits substantially faster training than the original CL4PHI (2023) implementation under identical hardware settings. This improvement arises from differences in objective formulation and optimization, rather than from margin-based learning being inherently inefficient. The CL4PHI implementation optimizes margin-based losses over individual phage–host pairs, requiring repeated operations over explicitly enumerated negative hosts. In contrast, CE4PHI adopts an InfoNCE-style cross-entropy formulation that jointly processes all candidate hosts for each phage, enabling efficient matrix-based optimization. As a result, CE4PHI achieves dramatic speedups of around 26.37 × on the CHERRY dataset and around 9.25 × on the Hi-C dataset over 300 training epochs, while maintaining strong species-level Top-1 accuracy. This efficiency makes CE4PHI well suited for scalable training on large multi-host PHI datasets.

## 4 Discussion

In this work, we proposed CE4PHI, a unified phylogeny-aware cross-entropy framework that formulates phage–host interaction identification from a positive–unlabeled perspective within deep metric learning. It combines InfoNCE-style multi-positive supervision for known infecting hosts with phylogenetically guided probability mass reallocation over unlabeled hosts to provide structured soft negative supervision. Across reference and metagenomic datasets, this design yields higher multi-host accuracy and improved species-level resolution, with favorable generalization behavior under cross-dataset evaluation relative to margin-based contrastive learning.

Traditional k-mer–based host prediction methods perform competitively on metagenomic datasets, where viral contigs are short, fragmented, and noisy, and local compositional signals provide robust cues for host association. In such settings, short k-mer statistics can effectively capture host-specific sequence biases even in the absence of complete genomes. Our contrastive learning framework can be viewed as a natural extension of k-mer–based approaches that preserves their strengths while addressing these limitations. Specifically, we retain FCGR-derived k-mer representations as inputs, thereby inheriting the robustness of compositional statistics, while further introducing a nonlinear encoder trained with a contrastive objective to transform these features into a shared embedding space. This representation-learning step enables the model to capture higher-order sequence organization and host phylogenetic structure beyond raw k-mer overlap. As a result, the proposed framework achieves consistently competitive performance across both metagenomic and whole-genome datasets.

From a formal perspective, the CE4PHI objective is closely related to the InfoNCE loss commonly used in contrastive learning, as both formulations are based on a softmax normalization over pairwise similarity scores between a phage and its candidate hosts. In particular, the InfoNCE-style component enforces concentration of probability mass on the true host by maximizing its relative similarity, thereby encouraging separation between positive and negative samples in the embedding space. Beyond this basic contrastive mechanism, CE4PHI further introduces phylogeny-aware supervision that structures the distribution of probability mass among negative hosts. Rather than treating all non-positive hosts as independent and equally repulsive, the phylogenetic tree defines soft supervisory signals that guide how attention should be allocated across related host candidates. Importantly, this structure is still realized through the same softmax-based formulation, ensuring a unified probabilistic interpretation. Under this view, CE4PHI can be interpreted as a form of supervised attention learning. The phage embedding acts as a query, host embeddings correspond to keys, and the softmax-normalized similarity scores define attention weights over candidate hosts. The InfoNCE component constrains attention to focus on true hosts, while the phylogenetic supervision further shapes the attention distribution among negative hosts. Together, these components enable CE4PHI to learn a structured and well-calibrated attention mechanism for phage–host association modeling.

The current CE4PHI implementation is that it primarily relies on k-mer–derived representations, specifically FCGR features, as input to the encoder. While k-mer statistics are robust for fragmented metagenomic sequences, they may not fully capture higher-level biological signals, such as gene content, protein sequence and domain information, or long-range genomic organization. Importantly, this limitation arises from the choice of input features rather than from the CE4PHI framework itself. The proposed softmax-based, phylogeny-aware contrastive objective is agnostic to the form of input representation and can, in principle, be extended to incorporate gene-level, protein-level, or pretrained sequence representations, providing a general and extensible framework for future integration of richer biological features.

## 5 Conclusion

In this study, we propose CE4PHI, a phylogenetic tree–aware positive–unlabeled deep metric learning framework for phage–host interaction prediction. By explicitly modeling unlabeled host candidates under phylogenetic guidance, CE4PHI moves beyond conventional classification and margin-based contrastive approaches that treat all non-positive pairs as negatives. Methodologically, CE4PHI unifies contrastive learning and deep metric learning through a softmax-based objective that integrates supervised InfoNCE for positive pairs with phylogeny-aware cross-entropy for non-positive samples. This formulation can be interpreted as supervised attention learning over candidate hosts, encouraging representations that are both discriminative and aligned with host evolutionary structure. Experiments on the Cherry benchmark and Hi-C-derived metagenomic datasets demonstrate that CE4PHI improves multi-host prediction accuracy, enhances species-level resolution, and exhibits stronger cross-dataset generalization, while substantially accelerating training compared with margin-based methods.

## Code availability

Related code, dataset splits and trained models can be found at: https://github.com/yaozhong/CE4PHI.

## Acknowledgments

The computing resources were provided by Human Genome Center, the Institute of Medical Science, the University of Tokyo. We also thank Martial Marbouty for providing the metagenomic Hi-C data.

## Footnotes

1 https://github.com/yaozhong/CL4PHI

