## Supplementary for "Phylogenetic tree-aware positive-unlabeled deep metric learning for phage–host interaction identification"

Supplementary material

1. Constructing phylogenetic tree for bacterial hosts
   1.1 GTDB-tk install

*Python -m pip install gtdbtk*

1.2 Running GTDB

Hosts may contain both archaea and hosts, run with the ref of bac120 or arc122 separately and then merge the distance matrix (e.g., 223 hosts of CHERRY dataset).

/mnt/ws1/phage_host2/data/hiC/host/merged_host

*gtdbtk de_novo_wf --genome_dir split/ --out_dir gtdb_output_arc/ --archaea --outgroup_taxon p__Bacillota --cpus 16 --extension fasta --tmpdir /mnt/ws1/tmp --skip_gtdb_refs*

*gtdbtk de_novo_wf --genome_dir split/ --out_dir gtdb_output/ --bacteria --outgroup_taxon p__Thermoproteota --cpus 16 --extension fasta --tmpdir /mnt/ws1/tmp*

***Note that the GTDBtk uses fastree for generating the nwk file, and the default setting builds ref and user provided fasta file. This requires large computational resources and prompt errors. The following solution can be used.

*export OMP_NUM_THREADS=8*

*export OMP_STACKSIZE=1G*

*ulimit -s unlimited*

*gzip -dc hiC_gtdb_bac_output/align/gtdbtk.bac120.user_msa.fasta.gz | FastTreeMP -wag -gamma -nosupport > hiC_gtdb_bac_output/align/gtdbtk.bac120.user.nwk*

2. HiC Data

| **phage_score** | **Count** | **Single host** | **Multi-host** | **Multi-host Percentage** |
| --- | --- | --- | --- | --- |
| Complete | 145 | 123 | 22 | 15.17 |
| High-quality | 937 | 760 | 177 | 18.89 |
| Medium-quality | 1801 | 1513 | 288 | 15.99 |
| Total | 2883 | 2396 | 487 |  |

| **phage_score** | **#host** | **Phage count** | **Multi-host percentage (%)** |
| --- | --- | --- | --- |
| **Complete** | 2 | 18 | 81.82 |
|  | 3 | 3 | 13.64 |
|  | 6 | 1 | 4.55 |
| **High-quality** | 2 | 124 | 70.06 |
|  | 3 | 37 | 20.90 |
|  | 4 | 10 | 5.65 |
|  | 5 | 4 | 2.26 |
|  | 6 | 1 | 0.56 |
|  | 7 | 1 | 0.56 |
| **Medium-quality** | 2 | 189 | 65.62 |
|  | 3 | 57 | 19.79 |
|  | 4 | 25 | 8.68 |
|  | 5 | 6 | 2.08 |
|  | 6 | 7 | 2.43 |
|  | 7 | 1 | 0.35 |
|  | 8 | 3 | 1.04 |

**249 hosts with species level taxnonomy (52 unique species)**

*s__Sutterella_wadsworthensis, s__Bacteroides_stercoris, s__Campylobacter_upsaliensis, s__Bacteroides_vulgatus, s__Clostridium_perfringens, s__Lactococcus_garvieae, s__Lactobacillus_reuteri, s__Bacteroides_uniformis, s__Corynebacterium_glutamicum, s__Acinetobacter_lwoffii, s__Blautia_wexlerae, s__Enterococcus_faecium, s__Ruminococcus_gnavus, s__Klebsiella_pneumoniae, s__Proteus_mirabilis, s__Streptococcus_salivarius, s__Acidaminococcus_intestini, s__Bifidobacterium_longum, s__Bifidobacterium_adolescentis, s__Citrobacter_freundii, s__Bacteroides_caccae, s__Enterobacter_cloacae, s__Bacteroides_xylanisolvens, s__Dorea_longicatena, s__Bacteroides_fragilis, s__Bifidobacterium_dentium, s__Bacteroides_ovatus, s__Lactobacillus_ruminis, s__Bacteroides_thetaiotaomicron, s__Streptococcus_thermophilus, s__Lactococcus_lactis, s__Parabacteroides_merdae, s__Lactobacillus_rhamnosus, s__Enterococcus_faecalis, s__Veillonella_parvula, s__Bacteroides_eggerthii, s__Streptococcus_parasanguinis, s__Clostridium_bolteae, s__Lactobacillus_salivarius, s__Leuconostoc_citreum, s__Acetobacter_pasteurianus, s__Zymomonas_mobilis, s__Delftia_acidovorans, s__Lactobacillus_buchneri, s__Clostridium_clostridioforme, s__Salmonella_enterica, s__Klebsiella_oxytoca, s__Bacteroides_dorei, s__Alteromonas_macleodii, s__Acinetobacter_johnsonii, s__Laribacter_hongkongensis, s__Bifidobacterium_breve*

Phage total number: *2883*, statsitics, sequence length

| **Data set** | **#Sample** | **min** | **25%** | **50% (median)** | **75%** | **max** | **mean** | **std** |
| --- | --- | --- | --- | --- | --- | --- | --- | --- |
| HiC raw | 2883 | 2,634 | 26,705.5 | 37,407.0 | 53,580.0 | 948,135 | 49,934.62 | 51,138.53 |

*Comparision with the cherry dataset*

| **Dataset** | **#Seq** | **Min** | **25%** | **Median** | **75%** | **Max** | **Mean** | **Std** |
| --- | --- | --- | --- | --- | --- | --- | --- | --- |
| **CHERRY_train** | 1175 | 2578 | 38,371 | 47,342 | 74,458 | 497,513 | 67,215.88 | 54,222.75 |
| **CHERRY_test** | 634 | 4623 | 40,471.75 | 50,146 | 75,992.25 | 348,113 | 69,761.45 | 50,746.35 |
| **CHERRY_val** | 131 | 3412 | 39,538 | 50,755 | 88,946 | 221,828 | 69,771.89 | 50,439.94 |
| **Total** | 1922 |  |  |  |  |  |  |  |

*Final usable HiC PHIs with species level information*

| **Category** | **Medium-quality** | **High-quality** | **Complete** | **Total** |
| --- | --- | --- | --- | --- |
| **Count** | 243 | 136 | 27 | 406 |

|  | **Count** | **Min** | **25%** | **Median** | **75%** | **Max** | **Mean** | **Std** |
| --- | --- | --- | --- | --- | --- | --- | --- | --- |
| **Value** | 406 | 3085 | 25319 | 37565 | 52672.75 | 561,493 | 47270.8 | 47680.72 |

According to the mappable host species information

| **phage_score** | **Multi-host** | **Single-host** |
| --- | --- | --- |
| Medium-quality | 38 | 205 |
| High-quality | 8 | 128 |
| Complete | 0 | 27 |
| **Total** | 46 | **360** |

========== SUMMARY ==========

[split 1] single-train/val, multi-test

Train: 280, Val: 71, Test: 55

[split 2] stratified

Train: 259, Val: 65, Test: 82

==============================

3. Model hyperparameters setting

We select model hyper-parameters based on the validation for search key model parameter spaces of batch_size = {32, 64, 128}, lr={1e-3, 1e-4, 1e-5}.

Encoder Model Architectures

|  | **CL4PHI** | **CE4PHI** |
| --- | --- | --- |
| *CNN-Encoder* | FCGR (1 × H × W)  → Conv2D(64, k=K, s=2) + BN + ReLU  → Conv2D(128, k=K, s=2) + BN + ReLU  → MaxPool(2×2)  → Flatten  → FC(512)  → Embedding | FCGR (1 × H × W)  → Conv2D(64, k=K, s=2) + BN + ReLU  → Conv2D(128, k=K, s=2) + BN + ReLU  → MaxPool(2×2)  → Flatten  → FC(512)  **→ L2 Normalize**  → Embedding |
| *Model Paramters* | 2,764,928 | 2,764,928 |
| *Default*  *Distance* | Euclidean | Chord |
| *Hyper-parameters* | LR=1e-3, margin=1, Batch_size=32 | LR=1e-5, Batch_size=32  Temperature=0.07, ce=0.02 |

4. Tree distance analysis

Cherry data Intra-genus and extra-genus tree distance comparision

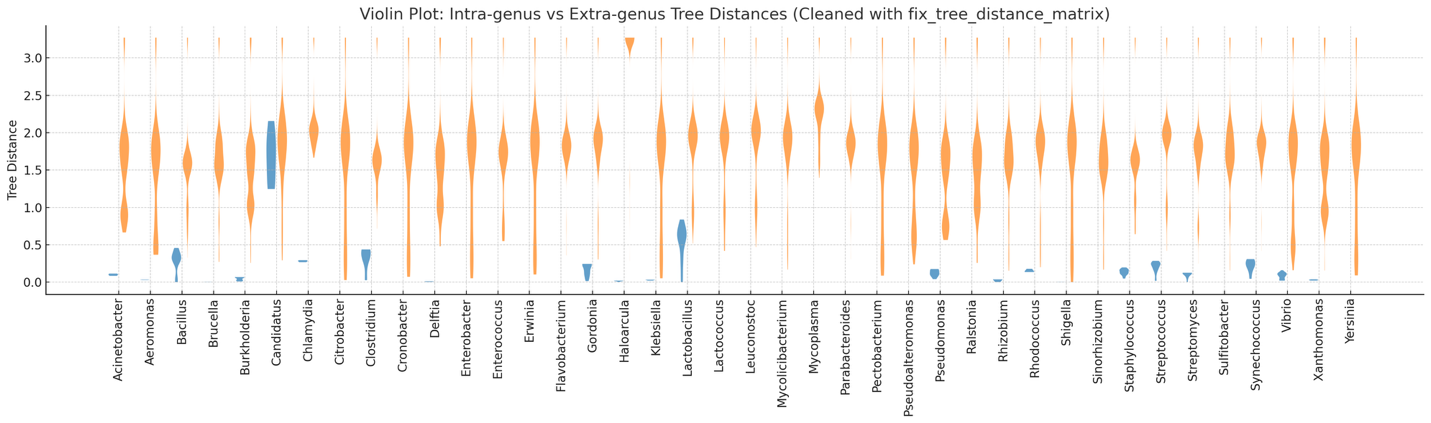

HiC data Intra-genus and extra-genus tree distance comparision

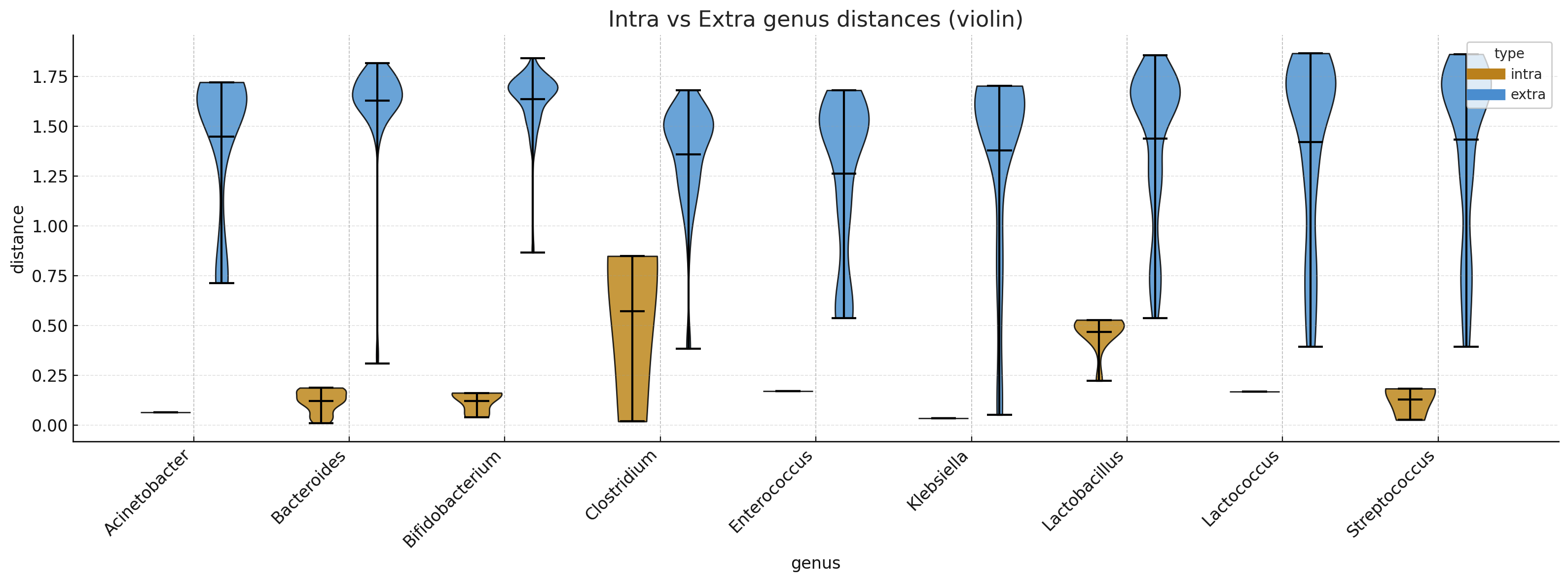

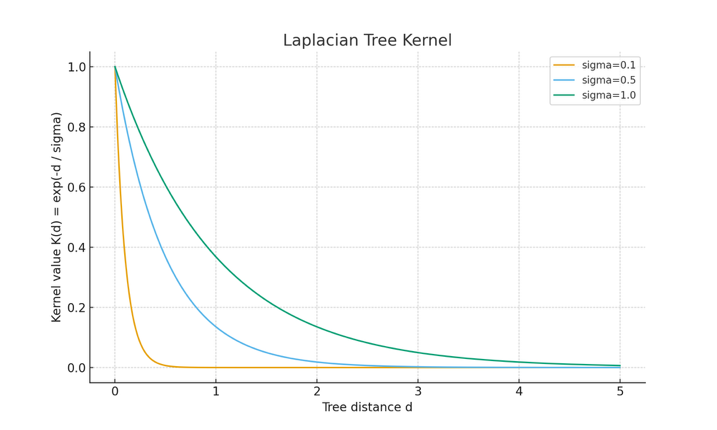

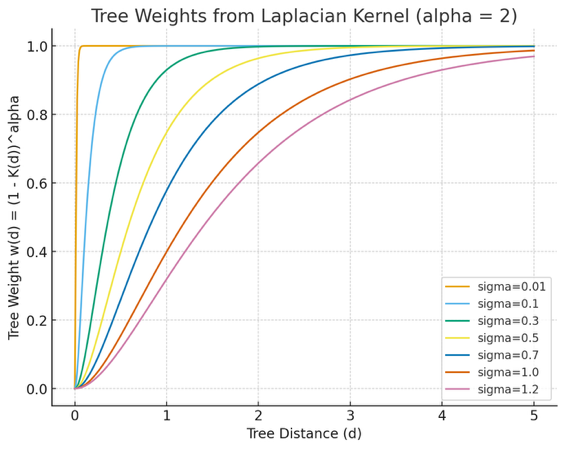
